# A synthesis of mapping experiments reveals extensive genomic structural diversity in the *Mimulus guttatus* species complex

**DOI:** 10.1101/330159

**Authors:** Lex Flagel, Benjamin K. Blackman, Lila Fishman, Patrick J. Monnahan, Andrea Sweigart, John K. Kelly

## Abstract

Understanding genomic structural variation such as inversions and translocations is a key challenge in evolutionary genetics. In this paper, we tackle this challenge by developing a novel statistical approach to comparative genetic mapping. The procedure couples a Hidden Markov Model with a Genetic Algorithm to detect large-scale structural variation using low-level sequencing data from multiple genetic mapping populations. We demonstrate the method using five distinct crosses within the flowering plant genus *Mimulus*. The synthesis of data from these experiments is first used to correct numerous errors (misplaced sequences) in the *M. guttatus* reference genome. Second, we confirm and/or detect eight large inversions polymorphic within the *M. guttatus* species complex. Finally, we show how this method can be applied in genomic scans to improve the accuracy and resolution of Quantitative Trait Locus (QTL) mapping.

**AUTHOR SUMMARY:** Genome sequences have proved to be a critical experimental resource for genetic research in many species. However, in some species there is considerable variation in genomic organization, making a single reference genome sequence inadequate. This variation can cause issues in interpreting genomic signals, such as those coming from trait mapping. We introduce a new statistical method and computational tools that use linkage information to reorganize a single reference genome to 1) repair genome assembly errors, and 2) identify variation between individuals or populations of the same species. Using this method we can create a new genome order that improves upon the reference genome. We apply this method to five crosses among plants in the *Mimulus guttatus* species complex. In this system we detect eight large chromosomal inversions and improve the resolution of a trait mapping study. This work highlights the utility of our method, and indicates how others studying diverse species might use them to improve their own research.

## INTRODUCTION

Over the last decade, genetic research has been revolutionized by the availability of whole genome sequences for many of the world’s medically, ecologically, and agriculturally important species. It has become increasingly clear that a single reference genome sequence is an insufficient description for many species. For example, a comparison of two maize accessions found that over 2,500 genes were present in only one of the two genomes [1]. Even in humans, a species with significantly less genetic diversity than maize, segregating structural and gene content polymorphisms are abundant [2]. Differences in gene copy number [3-6], variation in gene order [1, 7, 8] and chromosomal inversions [9-13] are not captured by a single reference genome, nor can they be annotated succinctly in relation to a single reference as is possible for Single Nucleotide Polymorphisms (SNPs). These structural and gene content variants have important phenotypic consequences in many species, highlighting the need for intensive study [14-18].

Recognizing structural variation is important for many of the experimental applications of genomic science. Consider trait-mapping analyses in species with segregating chromosomal variants. Many trait-mapping approaches (e.g. genome-wide association studies or bulked segregant analyses) rely on the accumulation of signals from adjacent genomic regions (windows) to establish significance. If gene order in the population under study differs from the reference genome, the proximity and presence of genomic windows can be incorrectly inferred. This, in turn, undermines interpretation of both the location and significance of QTLs [19]. Similar issues can arise in population genomic inferences, such as scans for selection or introgression [20, 21]. One solution is to make reference genomes for every divergent accession under study [4]. When this is not feasible, an alternative approach is to construct ‘pseudo-chromosome’ assemblies to better match structural variation in the focal accessions. Regardless, accounting for structural variation is an important challenge for the continued development of evolutionary genomics.

In this paper, we develop a new approach to pseudo-chromosome construction using comparative genetic mapping as the primary tool. In species with repeat rich genomes, whole genome shotgun sequencing and assembly typically yields many thousands of scaffolds. These scaffolds can be stitched together to form pseudo-chromosomes; often a necessary prerequisite to the trait-mapping and population genomic analyses. There are various techniques for making pseudo-chromosomes, such as following a BAC-tiling path [22], optical mapping using nanochannel arrays [23], or by localizing the scaffolds to markers on a genetic map [24, 25]. Genetic maps have proved to be invaluable tools for initial genome construction and pseudo-chromosome assembly [26-28]. We extend this approach using comparative genetic maps from five distinct crosses, allowing us to simultaneously improve the pseudo-chromosome representation of the reference genome and also identify large-scale variation in gene order, including chromosomal inversions and translocations.

Our approach utilizes data from low-coverage sequencing. RAD-seq [29, 30] and related reduced-representation methods [31-33] allow cost-effective genotyping of hundreds of recombinant individuals in species with limited molecular tools or known genetic markers. While RAD-seq data is often used to create *de novo* markers [34, 35], sequences can also be directly mapped to genomic scaffolds. Recombinant genotypes located to genomic scaffolds can then be used to assemble pseudo-chromosomes. Unfortunately, there are substantial challenges in constructing genetic maps from low-coverage sequencing data and also in the inference of map differences (i.e., the evidence for structural variation). New approaches are needed to address the following major methodological questions: What is the optimal means to convert sequence data into markers? How should we accommodate genotyping error in these markers given that the error rate is often high and variable among samples? After locating markers to genomic scaffolds, how do we obtain the optimal order and orientation of scaffolds into pseudo-chromosomes? Finally, how do we determine if putative differences between maps are real?

In this paper we develop a statistical procedure called **Genome Order Optimization by Genetic Algorithm** (GOOGA) that detects structural variation using marker data from multiple genetic mapping populations. Importantly for error-prone low-coverage genotyping, GOOGA propagates genotype uncertainty throughout the model, thus accommodating this source of uncertainty directly into the inference of structural variation. GOOGA couples a Hidden Markov Model (HMM) with a Genetic Algorithm (GA). The HMM yields the likelihood of a given ‘map’ (hereafter used to denote the ordering and orientation of scaffolds along a chromosome) conditional on the genotype data. The GA searches map space by creating new candidate orders which are recurrently fed to the HMM to diagnose their likelihood. Inference of recombination rate parameters and/or tests for differences in gene order are enabled by the fact that all calculations are conducted in the currency of likelihood.

As proof of concept, we apply GOOGA to RAD-seq data (MSG type [31, 36]) from five different mapping populations, each synthesized from a cross between lines within the *M. guttatus* species complex. The complex is a highly diverse clade of inter-fertile North American wildflowers [37-40]. The five mapping populations include both intra- and inter-specific crosses as well as multiple cross types (F2s, F3s, and RILs). Recombinant individuals from each mapping population were scored genome-wide for SNPs and then input to GOOGA. Starting from a rough-draft scaffold order [41], GOOGA produces an optimized ordering and orientation of genomic scaffolds for each population. From similarities across the assemblies of each mapping population, we are able to correct many errors (misplaced scaffolds) in the current reference genome of *M. guttatus*. Improved estimates for recombination rates indicate the effects of gene density and transposable elements on chromosomal variation in recombination. Comparisons among maps identify eight distinct structural polymorphisms, five of which were suggested by previous mapping studies [12, 13, 42-45]. Finally, we demonstrate how the application of GOOGA clarifies the results of a QTL mapping study by correcting errors in the reference genome.

## RESULTS

### Comparison of the *M. guttatus* V2 reference to the GOOGA optimized scaffold orders

The output of GOOGA is a chromosome-scale genetic map of scaffolds from the *M. guttatus* genome assembly, wherein we treat 100 kb intervals as distinct genetic markers. This map is an optimized pseudo-chromosome construction (all 14 chromosomes for all 5 mapping populations are reported in Supporting Table 1). The overall map lengths for the F2 crosses are 1278 cM for DUNTIL, 1523 cM for IMNAS, and 1258 cM for IMSWC. The average map length, 1353 cM, is shorter than previous F2 maps generated in *Mimulus* through PCR-based genotyping methods [12, 42, 45]. The IMF3 map is ≈50% longer than the F2 average (2043 cM), as expected given the extra generation of recombination between F2 and F3 generations. The IMPR map (1489 cM) is only slightly longer than the F2 average. There is additional recombination in the formation of the RILs, but the recombination parameter is specified differently (as related to crossover events) in the RIL HMM (see METHODS).

In aggregate, comparison of the GOOGA optimized maps from the five experiments to the *M. guttatus* V2 reference genome order (hereafter V2 map; [41]) indicates that a large number of updates to the reference genome are necessary. Although the maps differ importantly from each other, changes in scaffold order and orientation from the V2 map are usually shared among all five mapping populations. These regions likely reflect errors in genome assembly given that the reference genome was sequenced from an inbred line (IM62) used as one of the parents in two of our crosses (IMF3 and IMSWC). To illustrate this point, we compare the maximum likelihood of the IMF3 data under from the GOOGA optimized map to the V2 map (Figure 1) for three chromosomes. This intra-population cross is likely to be most congruent with the true order of the IM62 reference genome.

**Figure 1.**
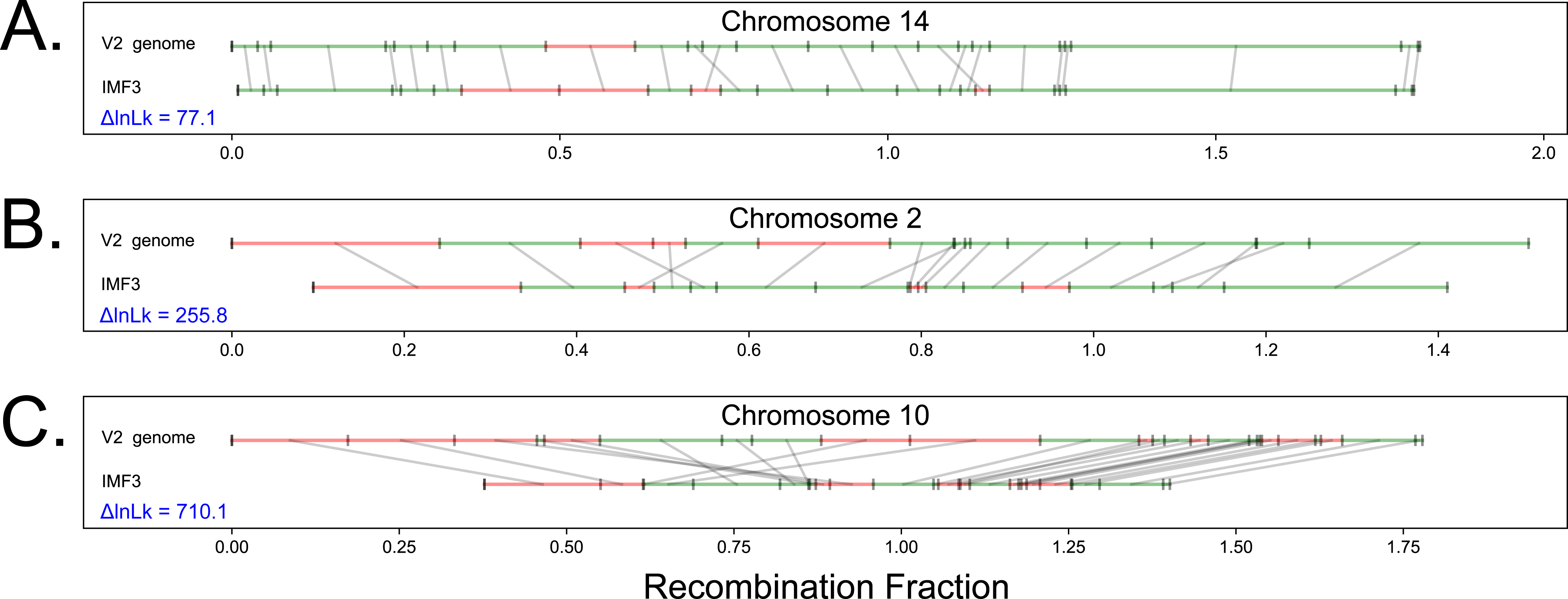
The improvement of the chromosome map between the *M. guttatus* V2 reference genome and the GOOGA optimized order for three chromosomes (IMF3 cross). Within each panel, the V2 order is shown above the IMF3 optimized order. Each genomic scaffold is drawn to its genetic map length (in Morgans) and denoted in green if it maps on the forward strand or red for the reverse strand. Grey lines connect the same scaffold in the V2 and optimized order.

The panels (Figure 1A-C) are ordered from most congruent with the V2 map (chromosome 14) to least congruent (chromosome 10), and the difference in log-likelihood (ΔlnLk) provides a measure of improvement in fit of the GOOGA relative to V2. ΔlnLk is computed by fitting the genotype data to both the GOOGA optimized and V2 maps, and then subtracting the former from the latter. In each case, recombination rates are estimated independently, and so ΔlnLk is determined entirely by differences in scaffold order and orientation. The maps for chromosome 14 (Figure 1A) are largely similar. However, the few differences, such as the changes in location and orientation of scaffolds 127, 211, 178, and 140b (inconsistency near the center of Figure 1A, Supporting Table 1), are sufficient for a large improvement in likelihood. The ΔlnLk of 77.1 (Supporting Table 2) corresponds to an improvement in likelihood greater than 10^33^, suggesting this new order fits the segregation data in the IMF3 population far better than the V2 map. Importantly, the updated ordering of scaffolds 127, 211, 178, and 140b is shared by the IMSWC, DUNTIL, and IMPR maps (Supporting Figure 1 and Supporting Table 1). The IMNAS map retains the [211,127] ordering of the V2 map, albeit with a flip of 127. However, this difference is not biologically compelling because there is no evidence of recombination in this region among the 91 F2s genotyped in the IMNAS population. Genome assembly by genetic mapping will often fail when there is no recombination to provide signal, a factor that must be taken into account when comparing maps.

Chromosome 2 (Figure 1B) is typical of most IMF3 contrasts. There are numerous rearrangements (e.g. the IMF3 sequence [44a, 212, 249] is inverted in the V2 map) and several scaffolds are flipped in place (including scaffold 81 that flanks [44a, 212, 249]). Apart from the increase in likelihood from the V2 map to the GOOGA optimized order (ΔlnLk = 255.8), it is noteworthy that the genetic length of chromosome 2 shrinks by ≈15% from V2 to GOOGA. This is because maximum likelihood of the rate parameters will compensate for bad joins by increasing recombination fraction (r) values. This effect is even more pronounced for chromosome 10 (Figure 1C), where there is a large increase in ΔlnLk. Among 70 chromosomes (5 crosses × 14 chromosomes), 23 (33%) chromosomes had ΔlnLK improvements greater than 500, while only 5 (7.1%) improved less than 100 (Supporting Table 2). The most significant alterations to the V2 map are in genomic regions harboring inversions, particularly on chromosomes 5, 8, and 10 (Figure 1C, Supporting Figure 1, and Supporting Table 2). The V2 map is based partly on genetic mapping data with segregation data from approximately 70 recombinant inbred lines from a cross between IM62 and DUN10 (J. Willis pers. comm.). DUN10 is a parent in our DUNTIL cross [35] and the aggregate of evidence (see below) suggests that DUN10 has chromosomal inversions (relative to IM62) on each of these chromosomes.

### Comparison of GOOGA maps to one another reveals known and new structural diversity

Our five mapping populations contain one intra-population cross (IMF3), two inter-population crosses (IMPR, IMSWC), a close interspecific-cross (IMNAS), and a more distant interspecific cross (DUNTIL). We observe structural polymorphisms by aligning the five maps to each other by chromosome. Figure 2 compares the maps for chromosome 10 which had previously been shown to harbor an inversion in IMPR [43]. As described in METHODS, we broke scaffold 13 into 13a and 13b based on a preliminary analysis of the DUNTIL data. GOOGA reassembled 13a and 13b into a continuous sequence for the IMF3 cross, but not the other crosses (Figure 2). There is minimal recombination between 13a and 13b in IMPR, IMNAS, and IMSWC because 13b is flanked by a large block of markers with nearly complete recombination suppression. This suppressed region, which represents at least 4.5 Mb of DNA, is freely recombining in IMF3 and DUNTIL but with a perfect reversal of marker order/orientation between those two crosses. From this we infer that the IMF3 parents (IM62 and IM767) each have inversion karyotype “A”, the DUNTIL parents (LVR and DUN) each have karyotype “B”, and the other three crosses are heterokaryotypic (one parent A and one B) for this inversion. Noting that IM767 is A, recombination suppression in the IMPR suggests the other parent in this cross (Point Reyes) has orientation B. By similar reasoning, we can conclude that SF and SWC also have orientation B. The right half of chromosome 10 is largely collinear among all five crosses, indicating the inversion is the primary influence on chromosome-wide likelihood. The GOOGA lnLk of the IMF3 data is −3784. If the IMF3 data is forced into the optimized DUNTIL order, the lnLk drops to −4074 (ΔlnLk = 290; Supporting Table 3). This gives strong statistical support of the inversion between the A and B homokarytypic crosses. The effect is less pronounced in the heterokaryotypic crosses. For example, the ΔlnLks of the IMNAS data when forced into the DUNTIL and IMF3 maps are 59 and 124, respectively (Supporting Table 3). Thus, as expected, the recombination suppression in this heterokaryotypic cross results in relatively weak support for either a pure A or B inversion orientation.

**Figure 2.**
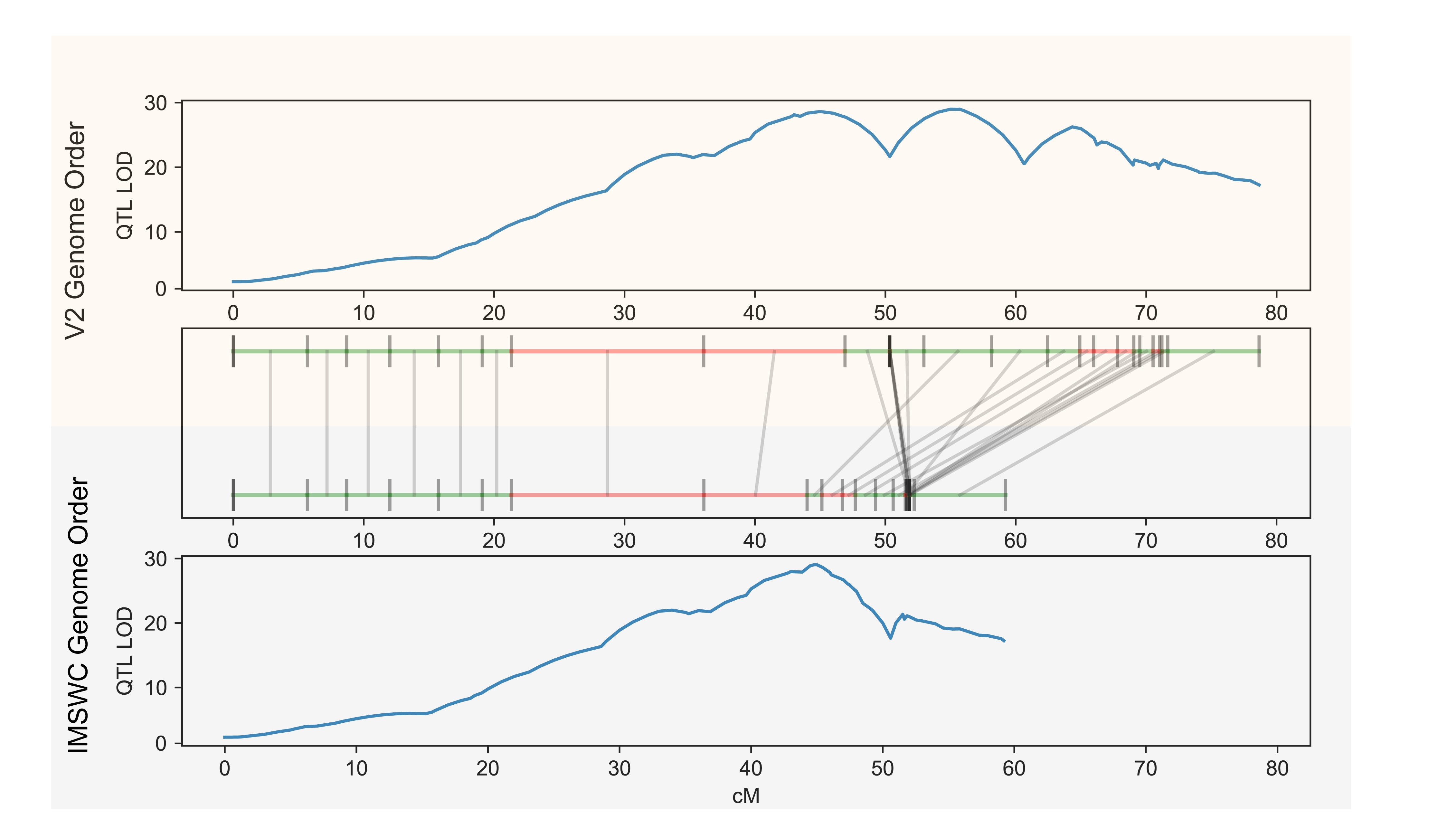
A reversal of genomic scaffolds due to an inversion is illustrated by the comparison of chromosome 10 maps for all five crosses. Each genomic scaffold is drawn to its genetic map length and denoted in green if it maps on the forward strand or red for the reverse strand. Grey lines connect the same scaffold between maps.

The inversion on chromosome 10 is the only case among these crosses where we see free recombination in both homokaryotypes and suppression in the heterokaryotypes. In all other cases, one or more crosses reveal recombination suppression, with at least one homokaryotypic cross also present among our five populations (Supporting Figure 1 and Supporting Table 1). Lowry and Willis [13] showed that reversal of marker order (as in Figure 2) for the inversion on chromosome 8. This feature is associated with annual versus perennial life-history within *M. guttatus*. Here, we see free recombination over the inverted region on chromosome 8 in IMF3, IMSWC, and IMNAS (annual × annual crosses), and suppression in IMPR and DUNTIL (annual × perennial *M. guttatus* and perennial *M. guttatus* × perennial *M. tilingii*, respectively). A similar pattern is noted for previously hypothesized inversions on chromosomes 5 (suppression in DUNTIL and IMPR) and 13 (suppression in DUNTIL) [44] and for the meiotic drive locus on chromosome 11 (suppression in IMF3) [46]. Given comparable evidence, we also identify three novel putative inversions (Supporting Figure 1 and Supporting Table 4). A region of at least 1.2 Mb spanning scaffolds 19, 73 and 65 on chromosome 2 is completely suppressed in the IMPR. There is substantial recombination across this region in other crosses: 8 cM in DUNTIL, 10 cM in IMF3, 7 cM in IMNAS, and 2 cM in IMSWC. A larger physical region (≈5 Mb on chromosome 7) is fully suppressed in IMPR but not the other crosses (20-45cM). Finally, a stretch of ≈4 Mb on chromosome 14 is suppressed in the IMSWC cross but not in other crosses (about 30 cM).

To provide a more general comparison of the extent of gene order differences among the crosses, we imposed the optimal map in every cross onto the genotypic data from every other cross and computed the ΔlnLk. Then we summed these for each chromosome. The larger the value of this sum, the greater degree of structural discrepancy we observe between the optimal fits of each cross (Table 1). As quantified by this metric, chromosome 10 has the greatest degree of structural discrepancy, an unsurprising result given the large polymorphic inversion (Figure 2). Chromosomes 5 and 11, which both show large tracts of reordered scaffolds and shared regions of recombination suppression among several crosses, rank the next highest in the degree of structural discrepancy among maps (Table 1). Surprisingly, the lowest value is for chromosome 8, which has a pairwise sum of ΔlnLk of only 75.4. The large inversion on chromosome 8 suppresses recombination in annual × perennial mapping populations, and as a consequence, the ordering of scaffolds within the inverted region is fairly arbitrary in those heterotypic crosses.

**Table 1:** Summed pairwise changes in log-likelihood values by chromosome given in descending order. For each chromosome we imposed the final scaffold order from every cross onto every other cross and calculated the difference in the natural log likelihoods (ΔlnLk) between the crosses own order versus this imposed order. Sums are given on the set of ΔlnLks for each chromosome. A large positive ΔlnLk indicates a pair of crosses with highly incompatible orders.

| Chromosome | Sum of Pairwise<br>$\Delta\ln Lk$ |
| --- | --- |
| 10 | 5581.1 |
| 5 | 5330.1 |
| 11 | 3503.0 |
| 3 | 2263.3 |
| 12 | 1449.2 |
| 13 | 1301.0 |
| 1 | 1203.7 |
| 7 | 845.8 |
| 6 | 828.2 |
| 2 | 782.0 |
| 4 | 610.8 |
| 14 | 551.0 |
| 9 | 442.1 |
| 8 | 75.4 |

The pairwise sum of ΔlnLks is determined by map changes, and these are rather few for chromosome 8. This result arises because the strongly supported, co-linear maps from the homokaryotypic crosses (IMF3, IMSWC, and IMNAS) are also largely reiterated in the suppressed crosses (DUNTIL and IMPR). However, all GOOGA maps of chromosome 8 represent a vast improvement over the V2 map (mean ΔlnLk vs V2 = 985, Supporting Table 2), suggesting the shared order that emerges from the inversion region is a large improvement over the reference genome.

### Correlation of estimated recombination rates with DNA composition

We next examined whether variation in the estimated recombination rates between successive markers within scaffolds correlated with aspects of the DNA content. We obtained an averaged recombination rate for each 200 kb window across mapping populations and then tested for association with the proportions of DNA annotated as coding sequence, transposable elements (TEs), and putative *M. guttatus* cent728 centromeric repeats [55] (Figure 3). Recombination rate is positively correlated with coding sequence density (Pearson’s r = 0.218) and is negatively correlated with TE density (Pearson’s r = −0.478). To test the impact of centromeric repeats [46] on recombination rate, we binned our 200 kb windows into those with < 5% centromeric repeat sequence vs. those with > 5%. Centromeres are expected to suppress recombination, and consistent with this prediction, we see a significant drop in recombination in windows with > 5% centromeric repeats (t-test p-value = 0.0003; Figure 3C). These results indicate that the recombination rates estimated by GOOGA fit well with biological expectations.

**Figure 3.**
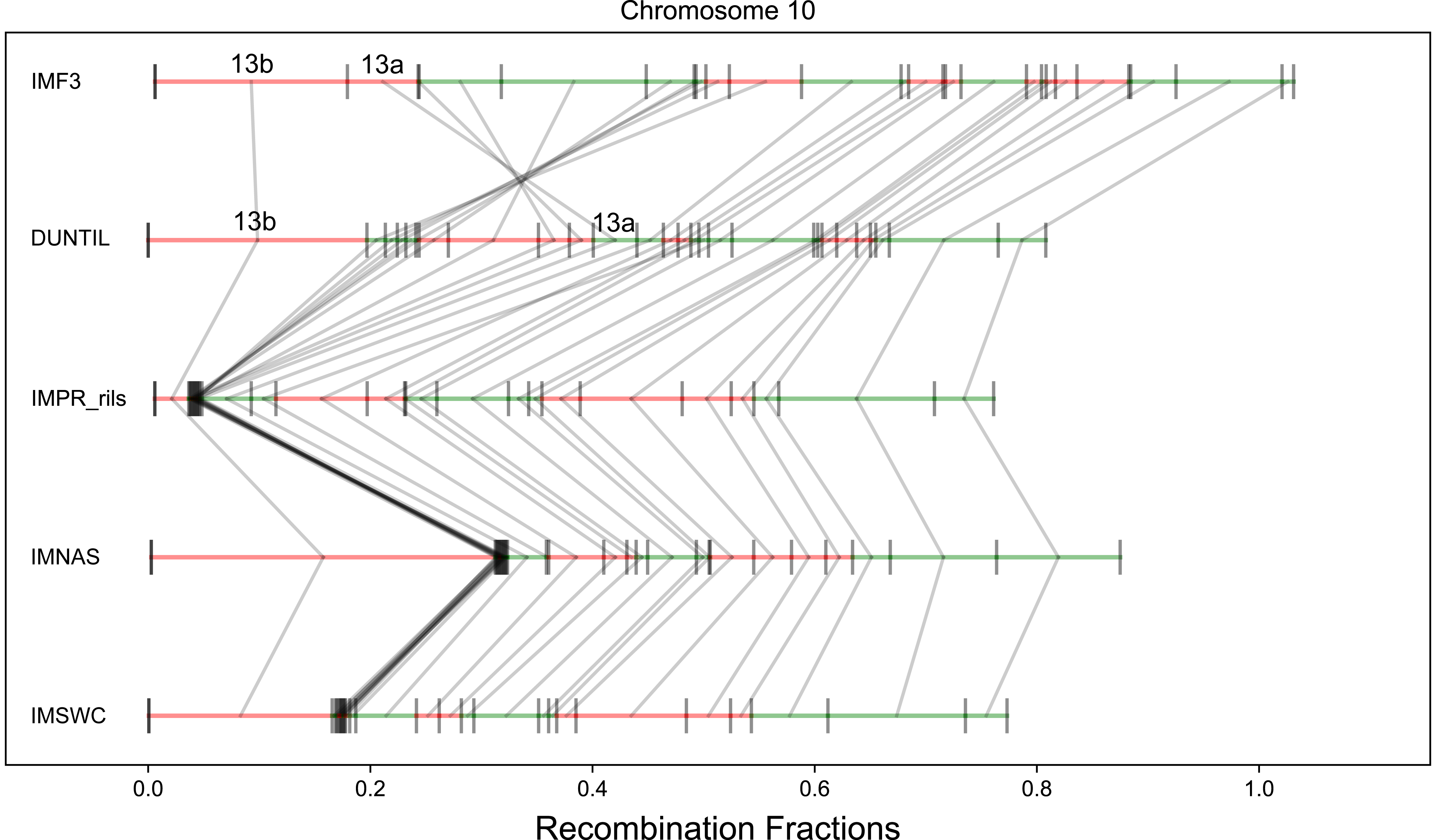
The effect of genomic features on recombination rate is illustrated. Panel A shows the positive correlation between the proportion of coding sequence and recombination rate. Panel B shows the negative correlation between the proportion annotated as a transposable element (TE) and recombination rate. For both panel A and B the red line markers the least squares best fit. Panel C shows the recombination rate distributions for 100 kb regions with >5% centromeric repeat content, or <5% centromeric repeat content.

## DISCUSSION

There has been a resurgence of interest among evolutionary biologists in structural variation, particularly in the contribution of chromosomal inversions to phenotypic variation, adaptive divergence among populations, and speciation [15-18, 47-51]. Inversions are routinely discovered from recombination suppression in genetic maps. They can be verified cytologically [11] or by reversal in marker order when comparing different genetic maps (e.g. [13], Figure 3). Our capacity to generate genetic maps has significantly advanced with RAD-seq and related genotyping platforms [29-33], particularly in non-model organisms. We developed GOOGA in response to these data, and with an eye toward statistically detecting structural variation. Accommodating error in map construction methodology is particularly important with low-coverage sequencing markers, which are abundant but error-prone. Propagating genotype uncertainty throughout the process to the assignment of map likelihoods provides a means to determine how strongly the underlying genotype data support apparent differences in scaffold order and orientation. The application of GOOGA to detect previously identified structural polymorphism in *Mimulus* illustrates how evidence of map differences manifest as differences in likelihood (Figures 1 & 2).

Andolfatto et al. [31] developed an HMM for use with RAD-Seq like data in genetic mapping. We have adopted the HMM approach here, although with numerous updates to both the model and implementation. First, we define markers within genomic windows, each inclusive of many SNPs, e.g. [43]. SNP calls from closely linked sites are aggregated to make a putative call for the ancestry of each genomic region (e.g. AA, AB, or BB) of each recombinant individual.

Given that the quality of individual DNA samples can vary greatly, we fit a model with individual specific genotyping error rates. The intervals between markers are treated differently depending on whether markers are within a genomic scaffold or between distinct scaffolds. The former are subject to tests for genotyping consistency given implied close linkage. The latter are explored using a genetic algorithm to order and orientation of scaffolds into chromosomes based on the likelihood of the data.

Window-based genotype calling is employed because there is substantial uncertainty associated with SNP genotyping from low-level data, particularly when reads are mapped to an unpolished draft genome. By applying calls to windows instead of individual SNPs, we sacrifice resolution to obtain more robust markers. In this application, we used 100 kb windows, which is about 0.3% of the average *Mimulus* chromosome. The resulting marker density is high relative to the number of recombination breakpoints per chromosome in F2, F3, and RIL individuals (typically 1-3; Supporting Table 1) and sufficient for testing alternative maps. However, scoring recombination events at the scale of 100-200 kb does limit inference of genomic features that determine recombination. Factors defined effectively at the 100 kb scale, such as the density of genes or transposable elements exhibit clear correlations with recombination, as expected from studies of other systems [52]. On the other hand, it is too coarse to evaluate finer scale determinants of recombination events. For example, population LD patterns suggest that recombination is strikingly elevated near the start site of genes in *M. guttatus* [41]. Figure 3A is fully consistent with this result: Mean recombination rate is correlated with the proportion of coding sequence, which is strongly correlated with number of gene start sites per window. However, at the resolution we selected, we cannot distinguish the effect of start sites from other features of gene rich regions.

The emission probabilities in the HMM portion of GOOGA are based on individualized genotyping error rates. This model feature stems from our observation that the quality of genotyping data varies quite substantially among samples, even of the same batch. This variability likely reflects stochastic factors, such as variation in the amount/quality of input DNA per individual. Regardless of cause, we find substantial differences in error rates among samples in all mapping populations (Supporting Table 6), justifying the need to account for these error rates explicitly and individually. While rates are typically low in absolute terms (medians around 0.01 for e_0i_ and e_2i_, much smaller for e_1i_; see METHODS), they are not negligible relative to actual recombination rates between adjacent markers. Differences in genotyping error rates provide key weights on the contributions of different individuals to the overall likelihood of a map and its associated collection of recombination rate estimates. A practical example of the utility of genotype filtering and genotyping error estimation is that it enabled the discovery of two novel putative inversions in the IMPR cross (on chromosomes 2 and 7). These were not identified in the original paper [43] because those authors imposed conservative thresholds both on marker inclusion and on whether individuals were included in the final map construction. Twice as many of the RIL plants are included in the present analysis and 25% more DNA is included in the map (1958 100 kb markers here as opposed to 3073 50 kb windows in [43]).

Admittedly, a considerable diversity of factors may complicate marker construction from low-coverage sequencing data [53-55]. We implement various filtering steps in GOOGA to mitigate these factors including SNP quality and allele frequency, SNP-level neighbor consistency tests, read-depth thresholds, marker-level (100 kb interval) neighbor consistency and heterozygosity tests, exclusion of individuals based on a high proportion of missing genotype calls, and/or high genotyping error rates. The goal of this filtering is to produce a marker set that is consistent with Mendelian segregation, but the filters will not always succeed. For example, a small region on chromosome 5 of the IMF3 cross involving only 7 markers (corresponding to the small scaffolds 226, 252, 94, 368 and 358) contributes 97 cM to the map length (over 40% of the total for this chromosome). It is possible that this a high recombination region (an unknown amount of DNA resides between these scaffolds), but it seems more likely a spurious inflation. These scaffolds map inconsistently in the other crosses – if they appear at all, they are not always adjacent. Of course, incorrectly locating good markers can produce the same “map inflation” effect as properly locating misleading markers. Also, even with fully accurate genotyping, recombination based methods cannot resolve non- or low-recombining areas. Sequencing-based methods [56] may be required to identify structural variants in these situations.

***Mimulus results*—**A tangible product of the application of GOOGA to *Mimulus guttatus* is that we substantially revise the reference genome of this species. The reference line (IM62) is used in two of our crosses, most importantly in IMF3 where it is crossed to another line from the same population (IM767). Excepting the meiotic drive locus on chromosome 11 [46], IM767 appears to be largely collinear with IM62 (the two lines have the same orientation at other putative inversions). Despite this, the GOOGA realignment of scaffolds yields a dramatic increase in IMF3 likelihood over the V2 assembly: ΔlnLk is 5464 when summed over all chromosomes (Figure 1 and Supporting Table 2). Map revisions are found on each chromosome, and supported not only by ΔlnLk within IMF3, but also the maps from the other crosses. Incomplete assembly is a common and important problem in genomics, particularly in species with complex patterns of repeats. The promising implication of Figure 1 is that the rough genome assembly of many species can be dramatically improved with a low-coverage sequencing of a mapping population. Moreover, GOOGA quantifies the magnitude of improvements in terms of increase in likelihood.

The alignment of maps for chromosome 10 illustrates the effect of an inversion. A 5 mb region (left portion of Figure 2) was previously identified as a putative inversion from recombination suppression in the IMPR [43]. This is clearly confirmed here by the inclusion of both homokaryotic crosses (A×A and B×B). The resulting ‘map flip’ (top two panels in Figure 2) effectively ascertains the scaffolds included within the inversion and their ordering. This example also highlights the importance of marker construction. While it is possible to construct markers *de novo* with RAD-seq data [35], here we delineate markers on a previously assembled set of reference DNA sequences (the genomic scaffolds from the *M. guttatus* reference genome) obtained from a specific source (sequencing of the IM62 inbred line). There are clear advantages to defining markers in this fashion, but care must be taken with this approach, especially in distantly related populations. For example, we initially assumed that the IM62 reference genome scaffolds correctly reflect the gene order for the other mapping populations. However, in our analysis of chromosome 10, we found it necessary to break scaffold 13 into two parts (13a and 13b), though it is continuous in IM62. GOOGA reannealed 13a and 13b in the IMF3 cross but inserted other scaffolds between them in other crosses due to a segregating inversion with a breakpoint contained within the original scaffold 13 (particularly DUNTIL; Figure 2). This implies that scaffold 13 was correctly assembled for IM62, but not for DUNTIL.

In the five crosses considered here, chromosome 10 is the only of the eight putative inversions where both homokaryotypic crosses are included. Reversal of marker ordering between homokaryotypic crosses was previously demonstrated for chromosome 8 [13], and with appropriate selection of parental lines, could likely be shown for others. However, the sort of reversal of genetic maps evident in Figure 2 requires polymorphism within both karyotypes. For several structural polymorphisms in *M. guttatus* [12, 57], the derived karyotype is essentially homogeneous (few or no SNPs). A cross between lines that share the derived karyotype can only identify recombination events (and thus order scaffolds) when polymorphisms exist in that cross. Although this will generally be the case for older inversions that have had time to accumulate mutational variants, this requirement may be limiting for ordering scaffolds within the derived allelic orientation of very young inversions.

The inclusion of phenotype data with the IMSWC cross illustrates the importance of scaffold ordering for downstream genetic analyses (Figure 4). Here, each F2 plant was scored for progression to flowering, a dichotomous trait in the experimental photoperiod, which was restrictive to floral induction for one cross parent. Application of the same QTL mapping procedure to the data produces radically different outcomes if markers are placed according to the current *M. guttatus* reference genome (V2 map is the top panel of Figure 4) or by the GOOGA optimized map (bottom panel of Figure 4). Using the latter (which has a ΔlnLk improvement of 1089 vs. V2; Supporting Table 2), the data suggest a single large-effect QTL localized to a map position 44-45 cM into chromosome 11, clearly upstream of the centromere (at approx. 52 cM). QTL mapping to the V2 orientation yields three distinct peaks, each with a high LOD score. The specific markers near the QTL peak in the GOOGA map are jumbled in the V2 map which splits the signal (the genotype-phenotype association) into three distinct parts. The V2 map is also “stretched” – expanded in recombination length by over 20 cM – likely to compensate for bad scaffold joins. Both of these effects are likely to impede QTL inference in places where the reference genome is misassembled.

**Figure 4.**
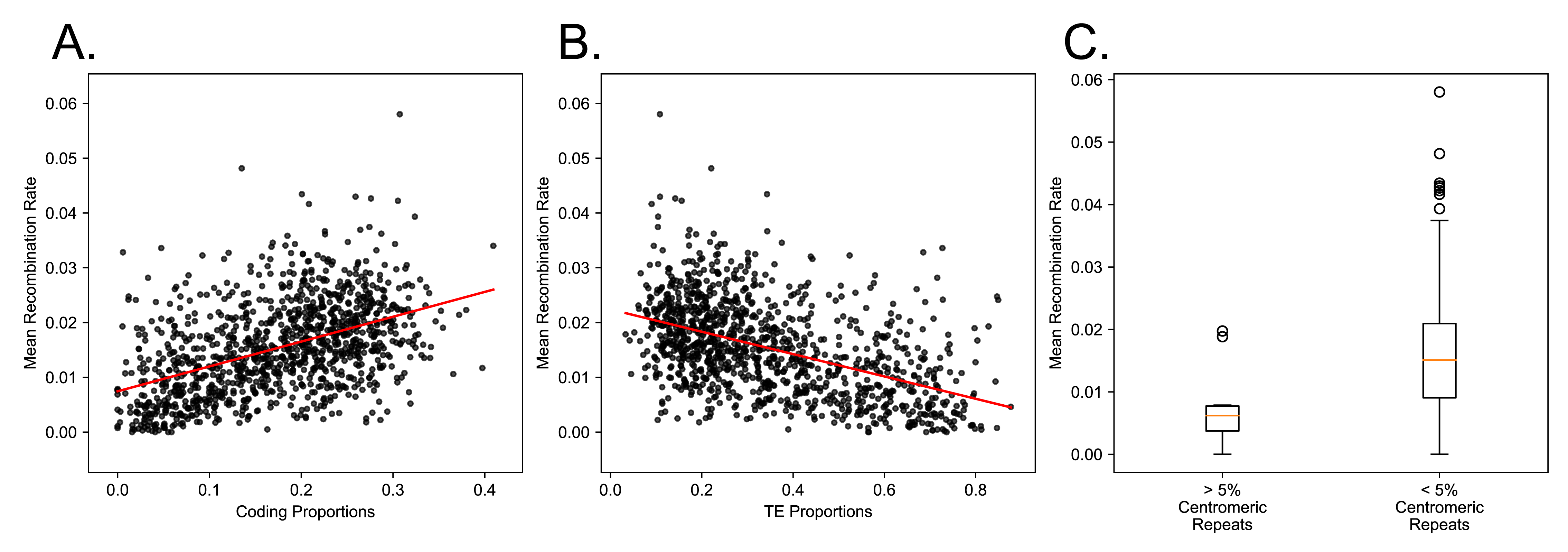
The results of QTL estimation using the V2 map (top panel) versus the IMSWC optimized map from GOOGA (bottom panel). The maps are aligned in middle panel.

Figure 4 suggests that GOOGA could be applied to test whether the reference genome order is consistent with that of the focal population in a population genetic scan or trait mapping experiment. Like *Mimulus*, many species and species complexes harbor significant segregating inversions and other gene order polymorphisms (e.g. *Drosophila melanogaster*, *Zea mays*, and the *Anopheles* and *Helianthus* species complexes [58-61]), but are represented by one or a small number of reference genomes. One could start from predefined genomic scaffolds, as we have done here, or by breaking the genome into small ‘pseudo-scaffolds’. Then GOOGA could be run on this reference genome with an appropriate mapping population. Using metrics like ΔlnLk, one could either confirm that the reference genome order is appropriate, or identify and fix issues as we have done in IMSWC for Figure 4. This approach may offer a compelling solution for species with incomplete genome assemblies such as *Mimulus*. Even in species with high-quality reference assemblies (e.g. *D. melanogaster*), this method could extend the utility of existing genomic resources in populations that are structurally diverged from the reference genome. This application would be particularly convenient in cases where trait mapping is being performed via RAD-Seq genotyping, as no additional data would need to be generated.

**Application—** The GOOGA pipeline is a set of modules including: (A) procedures to make genetic markers from low-coverage sequencing data in conjunction with a collection of genomic scaffolds, (B) a method to estimate genotyping error rates specific to each individual, (C) an application of the HMM to estimate recombination rates and obtain a likelihood for a specific ordering and orientation of genomic scaffolds, i.e. the map, and (D) a Genetic Algorithm (GA) to search map space to obtain pseudo-chromosomes that maximize the likelihood of the data. While GOOGA was developed as an integrated series of steps, the features and requirements of other datasets will differ from these *Mimulus* crosses. In these situations, one or more of the components might be used apart from the rest. We outline a few options below that could be appropriate for different species or scenarios.

The simplest application: Use (A) to create markers and then apply standard map making software [62, 63] to the resulting genotype matrix. If applied to the *Mimulus* mapping populations (or comparable datasets), this matrix directly from (A) contains a great excess of missing data. Extensive culling, both of individuals scored for too few markers and markers scored for too few individuals (e.g. [43]), is required to successfully apply standard map making programs. An alternative is to apply (A)-(B)-(C) to generate genetic markers. Assuming that the genomic scaffolds are generally reliable, the HMM will leverage data from neighboring windows to inform genotype calls. After obtaining the MLE on rates, one can extract posterior probabilities on genotypes and then impose ‘hard calls’, e.g. [36], to create a genotype matrix. We found this approach to be useful across a wide range of RAD-Seq coverages (see METHODS), including as little as ≈10 informative reads per windows. Another possibility is to replace the front end of the pipeline. If one has high certainty in the validity of individual SNPs (their location and scoring), it is natural to replace windows (A) with individual SNPs as the observed states of the HMM ([31], Figure 5). Finally, while we found the GA effective for searching map space (D), other map optimization methods exist (e.g., [64, 65]), and these alternative procedures may prove useful in searching order/orientation possibilities given that each can be assigned a likelihood.

**Figure 5.**
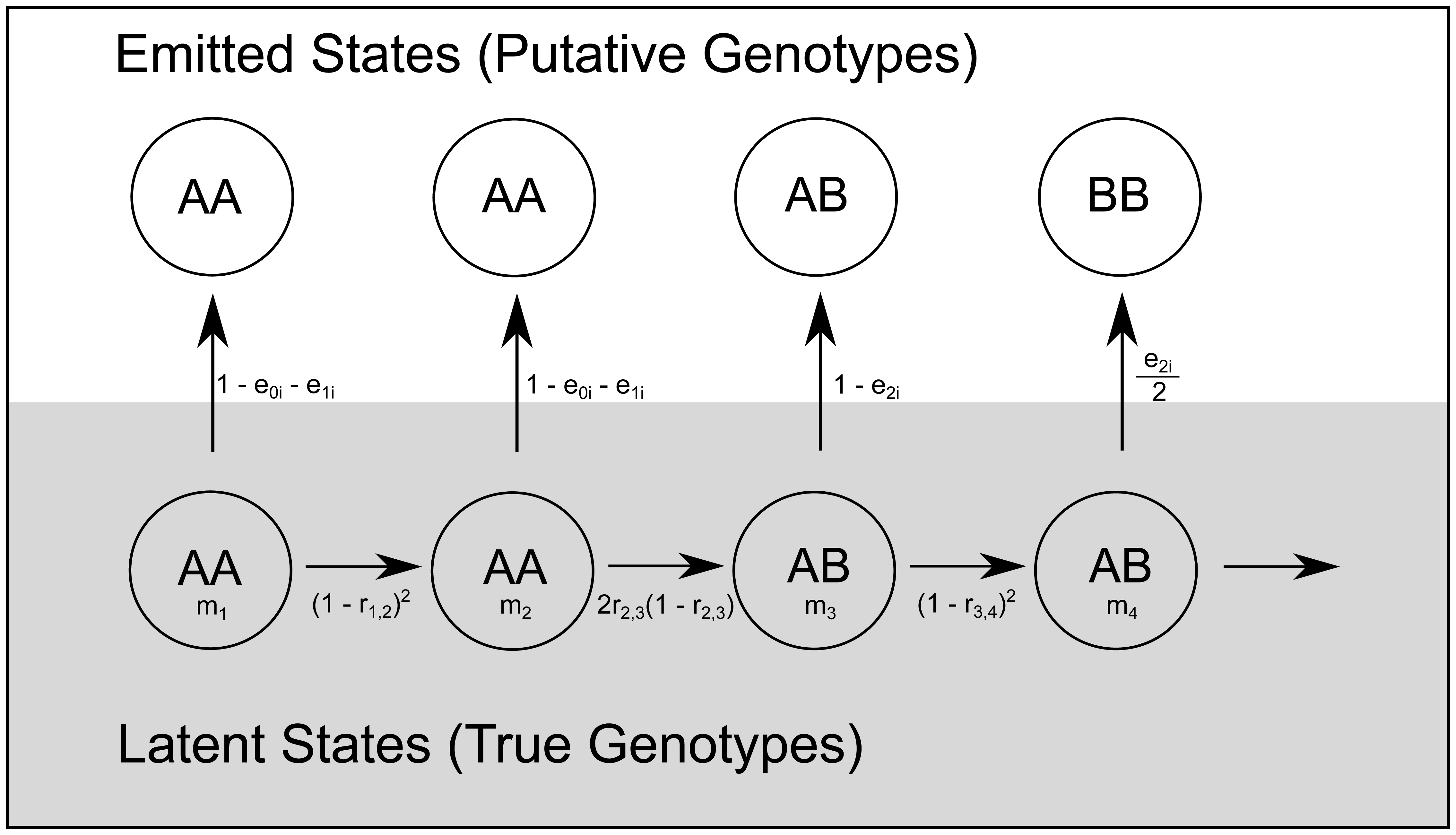
The structure of the Hidden Markov Model is illustrated for F2 individuals. Transitions between genotype states (AA, AB, or BB) for markers m_1_ through m_4_ in the latent state layer are determined by recombination fractions (r) between pairs of markers. The probabilities of each latent state estimate are then propagated into the emitted state layer with a genotyping error rate term that is specific to each individual (subscript i).

## MATERIALS AND METHODS

***Genomic scaffolds*—**The first iteration of the *M. guttatus* reference genome (v1, available at https://phytozome.jgi.doe.gov; login required), obtained from sequencing of a single inbred line (IM62; Iron Mountain, OR, USA), consists of over 2000 scaffolds. The longest are greater than 4 Mb in length (about 15% of an average *M. guttatus* chromosome) but the great majority of sequence is contained in scaffolds 10 kb − 1 Mb in size. The current assembly (V2 reference genome: available at https://phytozome.jgi.doe.gov; login required) orients most of the v1 scaffolds into chromosomal groups based on multiple sources of information [41]. For the present analysis, we revert back to the v1 scaffolds as a target for mapping sequence reads from recombinant individuals. We retain the set of breaks that were made to v1 scaffolds when creating the V2 build given that these breaks were largely corroborated by a subsequent mapping study [43]. We append a letter to names for broken v1 scaffolds, e.g. scaffold_97 is now scaffold_97a, scaffold_97b, and scaffold_97c. Next, we appended the mitochondrial and chloroplast genomic contigs to the scaffold list and masked repetitive regions to produce our read mapping target: Updated_v1_hardmasked.fa. The autosomal v1 scaffolds are the basic units for subsequent work. Markers are defined as contiguous stretches of DNA within these scaffolds.

***Creation and Genotyping of Mapping populations*—**Five crosses are analyzed in this study. Three crosses are F2s (IMNAS, IMSWC, DUNTIL), one is an F3 (IMF3), and one is a Recombinant Inbred Line panel (IMPR). The IMF3 population was founded by crossing two highly homozygous lines (IM62 and IM767) sampled from one population (Iron Mountain, OR, USA). A single F1 plant was selfed to create many F2s. F2 plants were randomly paired and crossed to produce F3 seed. Over 1000 F3 hybrids were grown to maturity and a random subset of these plants were genotyped for map construction. The IMSWC cross was formed from a cross between IM62 and SWC, an annual plant from a population of *M. guttatus* from Mapleton, OR, USA. The DUNTIL cross was formed by crossing an inbred line of *M. guttatus* (DUN10; Florence, OR, USA) to an inbred line of *Mimulus tilingii* (LVR; Inyo, CO, USA) [44]. For both IMSWC and DUNTIL, a single F1 plant was self-fertilized to create the F2 plants that were subsequently genotyped. IMNAS is a tri-parental cross: Two inbred lines from Iron Mountain (OR, USA) *M. guttatus* were each crossed to the SF5 (Sherars Falls, OR, USA) inbred line from selfing species *Mimulus nasutus*. The two interspecific F1s were then intercrossed [(SF5×IM160) × (SF5×IM767)] to produce the F2 plants. Because IM160 (like the reference line IM62) carries the driving *D* centromeric variant of LG11, which transmits nearly 100% via female function in heterozygotes with the *M. nasutus d* variant [46], this F2 segregates as a backcross in the D region. Finally, the IMPR are a set of highly homozygous lines derived from serially selfing progeny from a cross between IM767 and PR from the Point Reyes, CA, population of *M. guttatus* (see [66] for a description of RIL line formation).

The DNA extraction method and procedures for genotyping individuals using the Multiplexed-Shotgun-Genotyping (MSG; [31]) are described in Holeski et al [43]. Briefly, we digested DNA from each sample using a restriction enzyme (MseI or AseI) and then ligated unique Bar-Coded-Adapters (BCAs) to the resultant DNA fragments. After numerous cleaning steps, we size-selected our library for fragments between 250-425 bp. We then performed PCR reactions (14-18 cycles) using Phusion High-Fidelity PCR Master Mix and primers that bind to common regions in the BCAs. This elongates the molecules to contain necessary flanking sequence including the Illumina adaptors as well as additional indices to allow further multiplexing of samples within a single sequencing lane. Subsequent sequencing was performed using the Illumina instrument. MseI was used for the restriction digest to make a library for each the five mapping populations. We made a second library for the IMF3 population with a less frequent cutter, AseI. The genotype calls for IMF3 are synthesized from sequencing on these two libraries (see below). Details regarding the sequencing for DUNTIL [44], IMPR [43], IMNAS (Finseth et al, 2018, in prep) and IMSWC (Kooyers et al, 2018, in prep) are reported elsewhere, though briefly DUNTIL and IMSWC were sequenced at the Duke University Center for Genomic and Computational Biology, IMPR and IMF3 were sequenced at the University of Kansas Genome Sequencing Core, and IMNAS was sequenced at Hudson Alpha Genomic Services Laboratory.

Following sequencing, we demultiplexed reads into sample specific fastq files and then edited each with Scythe (https://github.com/vsbuffalo/scythe/) to remove adaptor contamination and then with Sickle (https://github.com/najoshi/sickle/) to trim low quality sequence. We used the *mem* function of BWA [67] to map read (or read pairs), one sample at a time, to Updated_v1_hardmasked.fa. Following read mapping, we identified putative SNPs using the UnifiedGenotyper function of the Genome Analysis ToolKit (GATK; [68]).

For each mapping population, the input data to create markers is a vcf file with all recombinant individuals scored for SNPs within individual reads or read pairs (RADtags) along with whole-genome sequencing data from one or both parental lines. We eliminated SNPs with a mapping quality score less than 30 or a minor allele frequency less than 0.1. We further thinned the data to a single SNP per RADtag, selecting the one with the most scored individuals. We then examined the relationship between read depth and apparent genotype for each SNP. There should be no relationship between the actual genotype and read depth, but MSG data exhibits two depth-related difficulties that require attention. First, there is typically under-calling of heterozygotes (relative to the binomial expectation) with low read depths [31]. This difficulty is addressed by the window based genotyping method (see below). Second, high read depth SNPs routinely deviate from Mendelian expectations, both in terms of allele frequency in the mapping population and in level of heterozygosity. After inspecting the distribution of reads called to each parent per individual for each possible read depth, we set a maximum median depth (across individuals scored for a SNP) specific to each mapping population: 10 for IMPR, 3 for IMSWC, 4 for DUNTIL, 10 for IMNAS, 5 for MseI libraries of IMF3, 20 for AseI libraries of IMF3. SNPs with excessive depth were suppressed.

Denoting the two parents of a mapping population as A or B, alleles were scored by assigning the reference base at each SNP to a specific parent (A or B). This was possible because one or both the parents in each crosses was MSG genotyped and/or fully genome sequenced. For the IMF3 population, one parent (IM62) is the reference genome sequence and the other (IM767) has been fully sequenced [37]. IM62 was also a parent in the IMSWC cross and the alternative base at a SNP is necessarily from the second parent (SWC). We used the IM767 genome sequence to polarize bases in the IMPR. For DUNTIL, we used genome sequences from both parents (LVR and DUN10) to specify base ancestry in the F2s [44]. For the IMNAS cross, we limited consideration to SNPs where an F1 hybrid between the IM767 and IM160 parents was homozygous for one base and the SF5 parent was homozygous for the alternative. After establishing calls to parent A/B for each SNP along each v1 scaffold, we imposed another layer of filtering based on consistency of the inferred parent within a recombinant individual along v1 scaffolds. We expect the same parentage of closely linked SNPs: If an individual is AA at a SNP, it should usually be AA at neighboring SNPs (excepting occasional recombination). SNPs exhibiting excessive disagreement were eliminated (details below).

***Window based genotype calling:*** We delineated markers as genomic windows of 100 kb in length within each v1 scaffold. The last marker on each scaffold included all remaining sequence beyond the last complete 100 kb segment, and was typically shorter. Within each window, we counted the number of reads scored as A and as B (across SNPs) within each recombinant individual. If the fraction of reads from parent A exceeded 95%, the individual was called AA for the marker; less than 5%, the genotype call was BB. If the fraction was between 25% and 75%, the individual was called AB. This method to assign putative genotypes (Figure 5) was designed to aggregate signal from closely linked SNPs that exhibit low average coverage and high variance in coverage among individuals. It compensates for heterozygote under-calling, where individual that are genuinely heterozygous appear homozygous at SNPs where the few reads present are all from one parent. Across a series of closely linked SNPs apparent homozygosity at individual sites may be high, but because alterative alleles (A or B) dropout at different SNPs, the similar overall frequencies of alternative bases within a window (summing across sites) indicated heterozygosity.

The genotype calling procedure defaults to ignorance (NN = No Call) if it cannot assign the marker to AA, AB, or BB following the rules above, or if fewer than 5 reads are mapped to the window. These cases of ambiguity can result either from mis-mapping of reads (in which case the read counts are misleading) or if recombination ‘splits’ the marker in an individual (in which case the true genotype is actually a combination of two different genotypes, e.g. AA-AB). Scoring either scenario as NN is suitable for downstream analysis by the HMM – data from neighboring markers will strongly inform inference of the underlying genotype at the NN marker.

The resulting genotype file for each individual is AA/AB/BB/NN at each marker. We imposed a third layer quality filtering at the scale of window-based genotypes. Each marker was scored for genotype frequencies and agreement of markers within 1 mb windows. We eliminated markers that were excessively heterozygous across mapping populations and/or exhibited high disagreement with neighboring markers. Within each mapping population, we suppressed loci that were excessively heterozygous in that population and/or had low numbers of called individuals. Finally, we imposed a cross-specific minimum number of called loci for a plant to be included in subsequent mapping. The programs used in the pipeline described above, as well as the likelihood algorithms described below, are available at (https://github.com/flag0010/GOOGA).

***Likelihood calculations*—**We treat recombination along each chromosome as a Markov Process with the true genotypes of each recombinant at each marker as the states of a Markov Chain. The likelihood takes the form of an HMM because the putative genotype calls (AA, AB, BB, or NN at each marker as obtained by methods described above) are treated as observed states [31, 69, 70] (Figure 5). The hidden states are the true underlying genotypes (AA, AB, or BB). The transition probabilities are contingent on recombination rates and the experimental design, the resulting HMM is thus non-homogeneous [71]. In an F2 population, the probabilities are (1-r)^2^, 2r(1-r), and r^2^ for AA transitioning to AA, AB, and BB, respectively [72]. The recombination rate, r, is specific to the flanking markers and we assume symmetry in transitions from BB to alternative genotypes. The transition probability of AB to AB is (1-r)^2^ + r^2^ and the probability AB to either homozygote is 2r(1-r). An additional round of recombination occurs in an F3 population in gamete formation by F2s. However, the probabilities have the same form except with an expected 50% increase in recombination rate values (assuming no change in crossover rates between F1s and F2s). In the RILs, heterozygosity has largely been eliminated by inbreeding. We suppress the remaining heterozygous regions by calling N at those loci. For the resultant genotypes, we stipulate the transitions probabilities as (1-r) and r for AA transitioning to AA and BB, respectively. For both RILs and F3s, the r parameter is actually a composite from multiple meioses. These apparent recombination rates are distinct from r of the F2 populations (the expected proportion of recombinant gametes from one round of meiosis) [73]. Thus, while the same markers are present across the different types of mapping populations, the absolute value of r will vary according to cross type. The emission probabilities are determined by individual-specific genotyping error rates (Figure 5). Three distinct error rates are estimated for each individual (i): the probability that a true homozygote yields a putative call to heterozygote (e_0i_) or to the opposite homozygote (e_1i_), and the probability that a heterozygote yields a call to one of the two homozygotes (e_2i_). Regarding the last rate, we assume that errors to either of the alternative homozygotes are equally likely (last emission in Figure 5).

The HMM is applied in three stages, first to estimate individual-specific error rates, then to obtain preliminary recombination estimates between markers within scaffolds, and finally to search for the optimal order and orientation of scaffolds within chromosomes (and also estimate recombination rates between scaffold ends within chromosomes). Genotyping error rates are estimated from transitions between genotypes *within* v1 scaffolds; we assume marker order in the scaffold is contiguous and correct. The transition probabilities depend on the true recombination rate per base pair, which is unknown and can vary across the genome. However, to estimate error rates, we fit a simple ‘homogeneous’ model assuming that recombination rate is proportional to physical distance between markers within scaffolds. We set rates to the genomic average of 5.0 cM/Mb for F2 populations and 10.0 cM/Mb for F3 and RIL populations. These point estimates are based on map lengths from numerous prior *Mimulus* studies [42, 45]. The likelihood of data from each scaffold of an individual plant is then a function of e_0i_, e_1i_, and e_2i_, and the likelihood for the entire plant is a product across scaffolds. For each plant we obtain the MLE of e_0i_, e_1i_, and e_2i_ via application of the forward-backward algorithm [74] coupled with the bfgs bounded optimization routine [75] of scipy.optimize (https://docs.scipy.org/doc/scipy/reference/optimize.html). We also used the bfgs optimizer to obtain MLE for recombination rates as described below.

The individual-specific error rates (reported as Supporting Table 6) are used in two ways. First, we used the estimates to cull plants with high error rates from subsequent analyses. For example, there were 130 F2s in the IMNAS population that passed preceding filters and fit for genotyping error rates. After excluding all plants where (e_0i_ + e_1i_ + e_2i_) ≥ 0.1, we obtained the set of 91 plants used for all downstream analyses. The post-error rates estimation sample sizes are 181 for IMF3, 872 for IMSWC, 205 for DUNTIL, and 260 for IMPR. Second, the genotyping error rates are treated as individual-specific constants in subsequent model fitting where marker-to-marker recombination rates (Figure 5) are free parameters. To obtain the MLE for recombination rates, we set the lower and upper bounds on rates as 0.0 and 0.2, respectively. A preliminary set of recombination estimates between markers within scaffolds was obtained by maximizing the likelihood with respect to r values for each scaffold across all plants within a mapping population. This collection of estimates is used in the GA search for optimal scaffold orders and orientations within linkage groups (described below), but not in the final maps. With one exception, the intra-scaffold r estimates are small, consistent with close linkage. We noticed a very high rate between two adjacent markers on scaffold 13 of the DUNTIL cross. The same interval exhibits normal (r < 0.01) recombination rates in other crosses. Given evidence for an inversion breakpoint (further evidence below), we split scaffold 13 into 13a and 13b 700 kb into the scaffold.

The third HMM application is to an entire linkage group for all individuals in a mapping population. This requires assignment of scaffolds to linkage groups. We tested assignments from the V2 build and found them consistent with our genotyping data (markers on the same linkage group exhibit positive association of genotypes). We were able to tie an additional 26 v1 scaffolds to linkage groups which were unassigned in the V2 build (Supporting Table 5). In the fully general model, there are L – 1 + K recombination rates to be estimated, where L is the number of scaffolds on the chromosome and K is the number of intra-scaffold rates within these scaffolds. The likelihood of a particular “map” (a specific ordering of scaffolds, each with a positive or negative orientation) is determined mainly by the data at the “joins” (where two scaffolds meet). In the GA runs that search map space (described below), we held intra-scaffold rates at their estimates from the stage 2 HMM. Thus, each evaluation was based on the likelihood of the map after optimizing relative to inter-scaffold rates. However, we apply the full model (all intra- and inter-scaffold recombination rates re-estimated) to our final map for each chromosome of each mapping population. In either case, we obtain the maximum likelihood via application of the forward-backward algorithm.

***The genetic algorithm*—**GOOGA maximizes the likelihood of the HMM using a genetic algorithm (GA) on a per chromosome, per mapping population basis. A GA is an algorithmic optimization scheme inspired by sexual reproduction and natural selection [76]. Below we use terms such as “individual”, “mutation”, “recombination”, and “selection” as they are frequently used in the GA literature, however, be aware we are not referring to biological entities or processes. To build the GA we first coded unique scaffold orders, including scaffold orientations (i.e. forward strand vs. reverse complement), to make an “individual”. Each individual represents a candidate solution among a population of individuals (*N*=19) competing against one another in a given generation. For each generation we used the HMM described above to calculate the likelihood of each individual. To preserve the best scaffold orders, we used a strategy called elitism (*E*), which allows a predetermined number (*E*=3 in our case) of the best individuals (i.e., highest likelihood scaffold orders) to go on to the next generation unchanged. To fill the remaining *N-E* spots in the next generation, we applied rank-based selection to select pairs of individuals to “mutate” and “recombine” into new individuals before adding to the next generation. To accomplish this, all *N* individuals were sorted in ascending order by likelihood and each individual (*i*) was assigned a rank (*R*_*i*_) of 1 to *N*. The probability of selecting an individual on the basis of its rank was 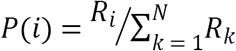. 16 pairs of unique individuals were randomly selected based on these probabilities, and subjected to mutation and recombination operations. We used a mutation scheme called “swap mutation”, where one individual was randomly selected from the pair, and then the location of up to four pairs of scaffolds within that individual were randomly swapped (while also flipping the orientation 50% of the time), thus creating a new “mutant” scaffold order. The mutated individual was then recombined with the other member of the pair. Recombination was performed using a scheme called the “order crossover”. This involves first choosing a random length segment of scaffolds from the donor individual and randomly inserting it into the scaffold order of the recipient individual to make a new individual. Each scaffold contributed by the donor is, now duplicated in the recipient individual. To fix this we deleted all the recipient individual’s copies of the duplicated scaffolds, while preserving the relative order of the non-duplicates. The swap mutation and ordered crossover do not mimic the mechanics of biological mutation or meiotic recombination. However both create diverse new individuals (i.e. scaffold orders) while preserving partial solutions from both the donor and recipient individuals. This allows the GA to build on past successes while exploring new scaffold orders.

The calculations were parallelized so that the HMM could be run simultaneously on each individual in the population. To further speed up the HMM, we implemented a memoization strategy that stored past recombination fraction (RF) results between scaffolds and injected those precomputed results into the future HMM calculations to avoid recalculation. To understand the mechanism of the memoization strategy, as a toy example and for the moment ignoring scaffold orientation, imagine a chromosome with four scaffolds A, B, C, and D and an initial individual ordered ABCD with RF rates computed between A-B, B-C, and C-D. After calculating the RF rates for this individual, we store the RF rate for the entire length (ABCD) and RF rates of all suborders of at least two scaffolds (ABC, BCD, AB, BC, CD), to create a catalog of previously computed rates. Then, when a future individual is produced, say DABC, we would search the catalog for matches starting with the longest members (ABCD) and then the next longest (ABC, BCD), and so on until the catalog is exhausted. For DABC, the first match would be the suborder ABC, which would supply RF rates for A-B and B-C, leaving D-A as the only missing RF rate to be calculated. After HMM calculation, all new suborders (DABC, DAB, and DA) would be added to the catalog for future use. This approach greatly speeds up calculation, especially at the later stages of optimization when many individuals tend to have large tracts of identical scaffold orders. We found that this approach reliably underestimates the likelihood value by a small amount when compared to the much slower processes of calculating all RF rates *de novo* for every new individual (among 750 random test samples all memoization estimates had a lower natural log-likelihood, with a median underestimate of 0.02%). This means the memoization method is consistently slightly conservative. For this reason, when a new individual was found to be among the elites (i.e. a promising new scaffold order), the program paused and recomputed this individual’s exact likelihood without precomputed RF rates and used this more precise likelihood for future ranking. The GA procedure is implemented in Python (version 2.7), utilizing functions from the scipy library (http://www.scipy.org/). The code is open source and available online (https://github.com/flag0010/GOOGA). All GA runs were initiated with a map based on the V2 genome order. The program was configured to terminate after 1000 generations with no change to the highest likelihood map or after running for 96 hrs.

After obtaining an optimal map for each cross considered in isolation, we contrasted the likelihood to that obtained under the V2 orientation. ΔlnLk is the difference in the natural log likelihood between two maps, each consisting of the same markers and their self-optimized recombination rates. We use this statistic as a measure of goodness of fit of data to alternative orders/orientations of scaffolds (e.g. Figure 1). We extracted MLE recombination rates from each mapping population to be compared to DNA level features such as amount of coding DNA, number of transposable elements, and the presence of centromeric DNA. For these analyses, we defined a 200 kb interval around each 100 kb-long marker, starting at the midpoint of the preceding marker and ending at the mid-point of the next marker. The analysis is defined at this scale to absorb recombination events that occur mid-marker (thus yielding NN at the focal site as described above). Consider the first three markers on a scaffold defined on the position ranges 0-100 kb, 100 kb-200 kb, and 200 kb-300 kb, respectively. We related the sequence interval from 50 kb-250 kb to the sum of the two rates (*r*_1,2_ + *r*_2,3_ of Figure 5). This analysis neglected very small scaffolds, the sequence at the ends of longer scaffolds, and the estimated rates between scaffolds (the amount and the features of the interceding DNA are unknown anyway). In this analysis, we also excluded regions where recombination is suppressed due to inversions (Supporting Table 4). We obtained a single rate for each interval by first standardizing map specific rates by the total length of each map and then calculated a weighted mean across populations. The weight given to estimates from each cross is proportional to the reciprocal of the genome-wide recombination rate variance: IMPR = 1, IMSWC =.899, IMNAS = 0.790, and DUNTIL = 0.677, and IMF3 = 0.380.

***QTL mapping in IMSWC population***: As a test case to demonstrate the value of applying GOOGA for genetic map construction prior to downstream applications, we compared quantitative trait locus (QTL) mapping results for the IMSWC cross using either the optimized scaffold order generated by our pipeline vs. the scaffold order of the M. guttatus v2 reference genome. For each of the 873 F2s used for genetic map construction, genotype posterior probabilities were emitted for each of the 111 markers defined for chromosome 11. This chromosome harbors a major QTL that contributes to variation in the ability to flower under 13 hour light: 11 hour dark conditions in this cross (Kooyers et al. 2018, in prep.). Genotypes were then assigned at each marker for each individual based on the genotype with a posterior probability > 0.95; otherwise, the genotype was called as missing. For each marker order, we calculated recombination frequencies (map function = haldane), imputed genotype probabilities at 1 cM steps (error probability = 0.001), and performed interval mapping using the binary model in R/QTL [48].

## ACKNOWLEDGEMENTS

The Minnesota Super Computing Institute and the Kansas University Center for Research Computing provided computing resources. We thank Brendan Epstein, Amanda Gorton, Joseph Guihlin, Thom Nelson, Courney Passow, Peter Tiffin, and Diana Trujillo for helpful comments on the manuscript.

## AUTHOR CONTRIBUTIONS

Conceived and designed the study: LEF, JKK. Performed the experiments: BKB, LF, PJM, AS, JKK. Contributed reagents/materials/analysis tools: LEF, BKB, LF, PJM, AS, JKK. Drafted the paper: LEF, JKK. All authors provided edits and approve the final manuscript.

## SUPPORTING INFORMATION LEGENDS

**Supporting Figure 1**: Comparative genetic maps for each of the five populations used in this study. For each chromosome the maps from each of the five populations are given. Each genomic scaffold is given in a separate color.

**Supporting Table 1**: Spreadsheet with marker orders. This spreadsheet includes the maps of every chromosome for each cross, and physical and genetic location of each marker. The last marker of each chromosome gives the map lnLk.

**Supporting Table 2**: Delta natural log likelihood (ΔlnLk) between GOOGA and V2 marker order. For each chromosome and each population the ΔlnLk is given. Large positive values indicate that the mapping data fits the GOOGA optimized order much better than the V2 genome order.

**Supporting Table 3**: Likelihood differences from imposing each GOOGA optimized map order onto every other population. All pairwise contrasts between population are given for each chromosome. “Home” is the focal data set, and “away” indicates the imposed map order. ll1 is the likelihood of “home” in it’s own map order, and ll2 is “home” in the “away” order. “Diff” is the difference between ll1 and ll2.

**Supporting Table 4**: All putative chromosomal inversions identified by GOOGA among the five mapping populations.

**Supporting Table 5**: Unplaced V2 genome scaffolds which were given a physical location from GOOGA. The table gives the v1 scaffold name and intervals for scaffolds that we were able to assign to one of the 14 *Mimulus* chromosomes. Their specific locations can be found in Supporting Table 1.

**Supporting Table 6**: Estimated genotype error rates for every individual from every population used in this study.

## REFERENCES

1. Hirsch CN, Hirsch CD, Brohammer AB, Bowman MJ, Soifer I, Barad O, et al. Draft assembly of elite inbred line PH207 provides insights into genomic and transcriptome diversity in maize. The Plant Cell. 2016;28(11):2700.

2. Sudmant PH, Rausch T, Gardner EJ, Handsaker RE, Abyzov A, Huddleston J, et al. An integrated map of structural variation in 2,504 human genomes. Nature. 2015;526(7571):75–81. doi: 10.1038/nature15394

3. Feulner PGD, Chain FJJ, Panchal M, Eizaguirre C, Kalbe M, Lenz TL, et al. Genome-wide patterns of standing genetic variation in a marine population of three-spined sticklebacks. Molecular Ecology. 2013;22(3):635–49. doi: 10.1111/j.1365-294X.2012.05680.x.

4. Long Q, Rabanal FA, Meng D, Huber CD, Farlow A, Platzer A, et al. Massive genomic variation and strong selection in Arabidopsis thaliana lines from Sweden. Nat Genet. 2013;45(8):884–90. doi: 10.1038/ng.2678

5. Springer NM, Ying K, Fu Y, Ji T, Yeh C-T, Jia Y, et al. Maize inbreds exhibit high levels of copy number variation (CNV) and presence/absence variation (PAV) in genome content. PLoS Genetics. 2009;5(11):e1000734. doi: 10.1371/journal.pgen.1000734.

6. Li Y-h, Zhou G, Ma J, Jiang W, Jin L-g, Zhang Z, et al. De novo assembly of soybean wild relatives for pan-genome analysis of diversity and agronomic traits. Nat Biotech. 2014;32(10):1045–52. doi: 10.1038/nbt.2979

7. Wright KM, Hellsten U, Xu C, Jeong AL, Sreedasyam A, Chapman JA, et al. Adaptation to heavy-metal contaminated environments proceeds via selection on pre-existing genetic variation. bioRxiv. 2015. doi: 10.1101/029900.

8. Naseeb S, Carter Z, Minnis D, Donaldson I, Zeef L, Delneri D. Widespread Impact of Chromosomal Inversions on Gene Expression Uncovers Robustness via Phenotypic Buffering. Molecular Biology and Evolution. 2016;33(7):1679–96. doi: 10.1093/molbev/msw045.

9. Coluzzi M, Sabatini A, della Torre A, Di Deco MA, Petrarca V. A Polytene Chromosome Analysis of the Anopheles gambiae Species Complex. Science. 2002;298(5597):1415–8. doi: 10.1126/science.1077769.

10. Fang Z, Pyhäjärvi T, Weber AL, Dawe RK, Glaubitz JC, Sánchez González JdJ, et al. Megabase-scale inversion polymorphism in the wild ancestor of maize. Genetics. 2012. doi: 10.1534/genetics.112.138578.

11. Krimbas CB, Powell JR. Drosophila Inversion Polymorphism. Boca Raton, FL: CRC Press; 1992.

12. Lee YW, Fishman L, Kelly JK, Willis JH. A Segregating Inversion Generates Fitness Variation in Yellow Monkeyflower (Mimulus guttatus). Genetics. 2016;202(4):1473–84. doi: 10.1534/genetics.115.183566.

13. Lowry DB, Willis JH. A Widespread Chromosomal Inversion Polymorphism Contributes to a Major Life-History Transition, Local Adaptation, and Reproductive Isolation. PLoS Biol. 2010;8(9):e1000500. doi: 10.1371/journal.pbio.1000500.

14. Stankiewicz P, Lupski JR. Structural variation in the human genome and its role in disease. Annual Review of Medicine. 2010;61(1):437–55. doi: 10.1146/annurev-med-100708-204735.

15. Schwander T, Libbrecht R, Keller L. Supergenes and Complex Phenotypes. Current Biology. 2014;24(7):R288–R94.

16. Love RR, Steele AM, Coulibaly MB, Traore SF, Emrich SJ, Fontaine MC, et al. Chromosomal inversions and ecotypic differentiation in Anopheles gambiae: the perspective from whole-genome sequencing. Molecular Ecology. 2016;25(23):5889–906. doi: 10.1111/mec.13888.

17. Dagilis AJ, Kirkpatrick M. Prezygotic isolation, mating preferences, and the evolution of chromosomal inversions. Evolution. 2016;70(7):1465–72. doi: 10.1111/evo.12954.

18. Samuk K. Inversions and the origin of behavioral differences in cod. Molecular Ecology. 2016;25(10):2111–3. doi: 10.1111/mec.13624.

19. Fransz P, Linc G, Lee C-R, Aflitos SA, Lasky JR, Toomajian C, et al. Molecular, genetic and evolutionary analysis of a paracentric inversion in *Arabidopsis thaliana*. The Plant Journal. 2016;88(2):159–78. doi: 10.1111/tpj.13262.

20. Hufford MB, Xu X, van Heerwaarden J, Pyhajarvi T, Chia J-M, Cartwright RA, et al. Comparative population genomics of maize domestication and improvement. Nat Genet. 2012;44(7):808–11. doi: 10.1038/ng.2309

21. Pyhäjärvi T, Hufford MB, Mezmouk S, Ross-Ibarra J. Complex Patterns of Local Adaptation in Teosinte. Genome Biology and Evolution. 2013;5(9):1594–609. doi: 10.1093/gbe/evt109.

22. Chen M, Presting G, Barbazuk WB, Goicoechea JL, Blackmon B, Fang G, et al. An Integrated Physical and Genetic Map of the Rice Genome. The Plant Cell. 2002;14(3):537–45. doi: 10.1105/tpc.010485.

23. Staňková H, Hastie AR, Chan S, Vrána J, Tulpová Z, Kubaláková M, et al. BioNano genome mapping of individual chromosomes supports physical mapping and sequence assembly in complex plant genomes. Plant Biotechnology Journal. 2016;14(7):1523–31. doi: 10.1111/pbi.12513.

24. Verde I, Jenkins J, Dondini L, Micali S, Pagliarani G, Vendramin E, et al. The Peach v2.0 release: high-resolution linkage mapping and deep resequencing improve chromosome-scale assembly and contiguity. BMC Genomics. 2017;18(1):225. doi: 10.1186/s12864-017-3606-9.

25. International Cassava Genetic Map Consortium (ICGMC). High-Resolution Linkage Map and Chromosome-Scale Genome Assembly for Cassava (*Manihot esculenta* Crantz) from 10 Populations. G3: Genes|Genomes|Genetics. 2015;5(1):133–44. doi: 10.1534/g3.114.015008.

26. Hahn MW, Zhang SV, Moyle LC. Sequencing, assembling, and correcting draft genomes using recombinant populations. G3. 2014;4(4):669–79. doi: 10.1534/g3.114.010264.

27. Mascher M, Muehlbauer GJ, Rokhsar DS, Chapman J, Schmutz J, Barry K, et al. Anchoring and ordering NGS contig assemblies by population sequencing (POPSEQ). The Plant Journal. 2013;76(4):718–27. doi: 10.1111/tpj.12319.

28. Nossa CW, Havlak P, Yue J-X, Lv J, Vincent KY, Brockmann HJ, et al. Joint assembly and genetic mapping of the Atlantic horseshoe crab genome reveals ancient whole genome duplication. GigaScience. 2014;3(1):1–21. doi: 10.1186/2047-217X-3-9.

29. Davey JW, Blaxter ML. RADSeq: next-generation population genetics. Briefings in Functional Genomics. 2010;9(5-6):416–23. doi: 10.1093/bfgp/elq031. PubMed PMID: PMC3080771.

30. Van Tassell CP, Smith TPL, Matukumalli LK, Taylor JF, Schnabel RD, Lawley CT, et al. SNP discovery and allele frequency estimation by deep sequencing of reduced representation libraries. Nat Meth. 2008;5(3):247–52. doi:

31. Andolfatto P, Davison D, Erezyilmaz D, Hu TT, Mast J, Sunayama-Morita T, et al. Multiplexed shotgun genotyping for rapid and efficient genetic mapping. Genome research. 2011;21(4):610–7. doi: 10.1101/gr.115402.110. PubMed PMID: WOS:000289067800011.

32. Elshire RJ, Glaubitz JC, Sun Q, Poland JA, Kawamoto K, Buckler ES, et al. A Robust, Simple Genotyping-by-Sequencing (GBS) Approach for High Diversity Species. PLoS ONE. 2011;6(5):e19379. doi: 10.1371/journal.pone.0019379.

33. Huang X, Feng Q, Qian Q, Zhao Q, Wang L, Wang A, et al. High-throughput genotyping by whole-genome resequencing. Genome Research. 2009;19(6):1068–76. doi: 10.1101/gr.089516.108.

34. Eaton DAR. PyRAD: assembly of de novo RADseq loci for phylogenetic analyses. Bioinformatics. 2014;30(13):1844–9. doi: 10.1093/bioinformatics/btu121.

35. Catchen J, Hohenlohe PA, Bassham S, Amores A, Cresko WA. Stacks: an analysis tool set for population genomics. Molecular ecology. 2013;22(11):3124–40. doi: 10.1111/mec.12354. PubMed PMID: PMC3936987.

36. Slotte T, Hazzouri KM, Stern D, Andolfatto P, Wright SI. GENETIC ARCHITECTURE AND ADAPTIVE SIGNIFICANCE OF THE SELFING SYNDROME IN CAPSELLA. Evolution. 2012;66(5):1360–74. doi: 10.1111/j.1558-5646.2011.01540.x. PubMed PMID: WOS:000303047300006.

37. Flagel LE, Willis JH, Vision TJ. The standing pool of genomic structural variation in a natural population of Mimulus guttatus. Genome biology and evolution. 2014;6(1):53–64.

38. Brandvain Y, Kenney AM, Flagel L, Coop G, Sweigart AL. Speciation and Introgression between *Mimulus nasutus* and *Mimulus guttatus*. PLoS Genetics. 2014;10(6):e1004410. doi: 10.1371/journal.pgen.1004410.

39. Puzey JR, Willis JH, Kelly JK. Population structure and local selection yield high genomic variation in Mimulus guttatus. Molecular Ecology. 2017;26(2):519–35. doi: 10.1111/mec.13922.

40. Vickery RK. Case studies in the evolution of species complexes in Mimulus. Evol Biol 1978;11:405–507.

41. Hellsten U, Wright KM, Jenkins J, Shu S, Yuan Y, Wessler SR, et al. Fine-scale variation in meiotic recombination in Mimulus inferred from population shotgun sequencing. Proc Natl Acad Sci. 2013;110. doi: 10.1073/pnas.1319032110.

42. Fishman L, Kelly AJ, Morgan E, Willis JH. A genetic map in the Mimulus guttatus species complex reveals transmission ratio distortion due to heterospecific interactions. Genetics. 2001;159(4):1701-16. PubMed PMID: PMC1461909.

43. Holeski L, Monnahan P, Koseva B, McCool N, Lindroth RL, Kelly JK. A High-Resolution Genetic Map of Yellow Monkeyflower Identifies Chemical Defense QTLs and Recombination Rate Variation. G3: Genes|Genomes|Genetics. 2014;4(5):813–21. doi: 10.1534/g3.113.010124.

44. Garner AG, Kenney AM, Fishman L, Sweigart AL. Genetic loci with parent-of-origin effects cause hybrid seed lethality in crosses between Mimulus species. New Phytologist. 2016;211(1):319–31. doi: 10.1111/nph.13897.

45. Fishman L, Willis JH, Wu CA, Lee YW. Comparative linkage maps suggest that fission, not polyploidy, underlies near-doubling of chromosome number within monkeyflowers (Mimulus; Phrymaceae). Heredity. 2014;112(5):562–8. doi: 10.1038/hdy.2013.143.

46. Fishman L, Saunders A. Centromere-Associated Female Meiotic Drive Entails Male Fitness Costs in Monkeyflowers. Science. 2008;322(5907):1559–62.

47. Tuttle Elaina M, Bergland Alan O, Korody Marisa L, Brewer Michael S, Newhouse Daniel J, Minx P, et al. Divergence and Functional Degradation of a Sex Chromosome-like Supergene. Current Biology. 2016;26(3):344–50. doi: https://doi.org/10.1016/j.cub.2015.11.069.

48. Feulner PGD, De-Kayne R. Genome evolution, structural rearrangements and speciation. Journal of Evolutionary Biology. 2017;30(8):1488–90. doi: 10.1111/jeb.13101.

49. Wellenreuther M, Rosenquist H, Jaksons P, Larson KW. Local adaptation along an environmental cline in a species with an inversion polymorphism. Journal of Evolutionary Biology. 2017;30(6):1068–77. doi: 10.1111/jeb.13064.

50. Kozak GM, Wadsworth CB, Kahne SC, Bogdanowicz SM, Harrison RG, Coates BS, et al. A combination of sexual and ecological divergence contributes to rearrangement spread during initial stages of speciation. Molecular Ecology. 2017;26(8):2331–47. doi: 10.1111/mec.14036.

51. Santos J, Pascual M, Fragata I, Simões P, Santos MA, Lima M, et al. Tracking changes in chromosomal arrangements and their genetic content during adaptation. Journal of Evolutionary Biology. 2016;29(6):1151–67. doi: 10.1111/jeb.12856.

52. Kent TV, Uzunović J, Wright SI. Coevolution between transposable elements and recombination. Philosophical Transactions of the Royal Society B: Biological Sciences. 2017;372(1736). doi: 10.1098/rstb.2016.0458.

53. Gautier M, Gharbi K, Cezard T, Foucaud J, Kerdelhué C, Pudlo P, et al. The effect of RAD allele dropout on the estimation of genetic variation within and between populations. Molecular Ecology. 2013;22(11):3165–78. doi: 10.1111/mec.12089.

54. Davey JW, Cezard T, Fuentes-Utrilla P, Eland C, Gharbi K, Blaxter ML. Special features of RAD Sequencing data: implications for genotyping. Molecular Ecology. 2013;22(11):3151–64. doi: 10.1111/mec.12084.

55. Monnahan PJ, Colicchio J, Kelly JK. A genomic selection component analysis characterizes migration-selection balance. Evolution. 2015;69(7):1713–27. doi: 10.1111/evo.12698.

56. Corbett-Detig RB, Cardeno C, Langley CH. Sequence-Based Detection and Breakpoint Assembly of Polymorphic Inversions. Genetics. 2012;192(1):131–7. doi: 10.1534/genetics.112.141622.

57. Fishman L, Kelly JK. Centromere-associated meiotic drive and female fitness variation in Mimulus. Evolution. 2015;69(5):1208–18. doi: 10.1111/evo.12661.

58. Corbett-Detig RB, Hartl DL. Population Genomics of Inversion Polymorphisms in Drosophila melanogaster. PLoS Genetics. 2012;8(12):e1003056. doi: 10.1371/journal.pgen.1003056.

59. Ayala D, Ullastres A, González J. Adaptation through chromosomal inversions in Anopheles. Frontiers in Genetics. 2014;5(129). doi: 10.3389/fgene.2014.00129.

60. Barb JG, Bowers JE, Renaut S, Rey JI, Knapp SJ, Rieseberg LH, et al. Chromosomal Evolution and Patterns of Introgression in *Helianthus*. Genetics. 2014;197(3):969–79. doi: 10.1534/genetics.114.165548.

61. Fang Z, Pyhäjärvi T, Weber AL, Dawe RK, Glaubitz JC, González JdJS, et al. Megabase-Scale Inversion Polymorphism in the Wild Ancestor of Maize. Genetics. 2012;191(3):883–94. doi: 10.1534/genetics.112.138578.

62. Arends D, Prins P, Jansen RC, Broman KW. R/qtl: high-throughput multiple QTL mapping. Bioinformatics. 2010;26(23):2990–2. doi: 10.1093/bioinformatics/btq565.

63. Iwata H, Ninomiya S. AntMap: Constructing genetic linkage maps using an ant colony optimization algorithm. Breeding science. 2006;56:371–7.

64. Green PJ. Reversible jump Markov chain Monte Carlo computation and Bayesian model determination. Biometrika. 1995;82(4):711–32. doi: 10.1093/biomet/82.4.711.

65. Tang J, Moret BME. Scaling up accurate phylogenetic reconstruction from gene-order data. Bioinformatics. 2003;(Eleventh International Conference on Intelligent Systems for Molecular Biology).

66. Holeski LM, Chase-Alone R, Kelly JK. The Genetics of Phenotypic Plasticity in Plant Defense: Trichome Production in Mimulus guttatus. American Naturalist. 2010;175(4):391–400. doi: 10.1086/651300. PubMed PMID: ISI:000275167100001.

67. Li H, Durbin R. Fast and accurate short read alignment with Burrows-Wheeler Transform. Bioinformatics. 2009;25:1754–60.

68. McKenna A, Hanna M, Banks E, Sivachenko A, Cibulskis K, Kernytsky A, et al. The Genome Analysis Toolkit: A MapReduce framework for analyzing next-generation DNA sequencing data. Genome Research. 2010;20(9):1297–303. doi: 10.1101/gr.107524.110.

69. Broman KW, Sen S. A Guide to QTL Mapping with R/qtl. New York.: Springer-Verlag; 2009.

70. King EG, Macdonald SJ, Long AD. Properties and Power of the Drosophila Synthetic Population Resource for the Routine Dissection of Complex Traits. Genetics. 2012;191(3):935–49. doi: 10.1534/genetics.112.138537.

71. Zamanzad GF, Jürgen C, Tomasz B. A Nonhomogeneous Hidden Markov Model for Gene Mapping Based on Next-Generation Sequencing Data. Journal of Computational Biology. 2015;22(2):178–88. doi: 10.1089/cmb.2014.0258. PubMed PMID: 25611462.

72. Fisher R, Balmukand B. The estimation of linkage from the offspring of selfed heterozygotes. Journal of Genetics. 1928;20(1):79–92. doi: doi:10.1007/BF02983317.

73. Martin OC, Hospital F. Two-and Three-Locus Tests for Linkage Analysis Using Recombinant Inbred Lines. Genetics. 2006;173(1):451–9. doi: 10.1534/genetics.105.047175.

74. Rabiner LR, Juang BH. An introduction to hidden Markov models. IEEE ASSP Magazine. 1986;January:4–15.

75. Byrd RH, Lu P, Nocedal J. A Limited Memory Algorithm for Bound Constrained Optimization. SIAM Journal on Scientific and Statistical Computing. 1995;16(5):1190–208.

76. Holland JH. Adaptation in natural and artificial systems: an introductory analysis with applications to biology, control, and artificial intelligence. Cambridge, Massachusetts: MIT Press; 1992.

